# Socially plastic responses in females are robust to evolutionary manipulations of adult sex ratio and adult nutrition

**DOI:** 10.1101/2023.09.28.559913

**Authors:** N McConnell, W Haerty, MJG Gage, T Chapman

**Affiliations:** School of Biological Sciences, University of East Anglia, Norwich Research Park, Norwich, NR4 7TJ, UK; The Earlham Institute, Norwich Research Park, Norwich, NR4 7UZ, UK

**Keywords:** sexual selection, plasticity, mating behaviour, experimental evolution

## Abstract

**Background:** Socially plastic behaviours are widespread among animals and can have a significant impact on fitness. Here we investigated whether the socially plastic responses of female *Drosophila melanogaster* can evolve in predictable ways following long term manipulation of adult sex ratio and adult nutrient availability. Previous reports show that female *D. melanogaster* respond plastically to their immediate same-sex social environment by altering their fecundity, laying fewer eggs after they mate if previously exposed to other females, Fecundity is also highly sensitive to a female’s immediate nutritional status, being significantly reduced when dietary protein in particular is scarce. On this basis, we predicted that an evolutionary history of exposure to variation in adult sex ratio and adult nutritional environment would select strongly upon a female’s plastic fecundity responses.

**Results:** We used females that had been drawn from replicated lines that had experienced an evolutionary history of male biased, female biased or equal adult sex ratios and either standard or low-quality adult nutrition. We tested the specific predictions that a history of elevated competition among females (in female-biased regimes) would select for increasingly sensitive plastic fecundity responses to the presence of conspecifics, and that these would be magnified under poor nutritional resource regimes. In contrast to the expectations, we found that the plastic responses in females were strikingly robust to perturbations of both sexual competition and nutrient availability and did not differ significantly across any of the evolutionary regimes. The lack of response is not explained by an insufficient strength of selection. For example, among females held in isolation prior to mating, we did observe significant evolutionary responses in virgin egg according to nutritional regime and in virgin egg retention to sex ratio regime.

**Conclusion:** The lack of variation in the existence and magnitude of predicted plasticity is consistent with the idea that the costs of maintaining fecundity plasticity in females are low, benefits high, and that plasticity itself can be hard wired.

## Background

Plastic responses expressed by individuals in response to environmental cues can be vital components of fitness across many different organisms [1-8]. Such responses allow organisms to match their reproductive effort or tailor their life history to the expected or prevailing environment, and thus optimise their fitness [9, 10]. Plastic responses can be manifested in many different ways. For example, they can be influenced by conditions experienced by parents, in anticipation of the environment likely to be experienced by offspring [3, 11]. They may also be set during development in anticipation for the expected adult environment [12, 13]. Behavioural plasticity or allocation of resouces to reproduction may also vary in response to the immediate conditions experienced during adulthood [1]. Plastic responses to environmental conditions such as diet can also balance lifespan and reproductive success according to the level of resources available [14-16] and can even change the sign of trade offs between lifespan and reproductive effort [17].

The capacity for individuals to express plasticity will be affected by the potential fitness gains of doing so balanced against any costs of being plastic [4-6, 8]. The extent, tempo and predictability of variation in expected or prevailing environments is expected to be a crucial determinant of the potential fitness benefits of plasticity. For example, if environments change very rapidly, plasticity responses may not easily be matched to them [18]. In contrast, if environments are generally stable, the fitness benefits of plasticity are expected to be minimal [19]. However, much about the evolutionary drivers, pace and extent of plasticity evolution remains unclear [20]. This is the topic we address here by testing key hypotheses for the evolution of reproductive plasticity in females drawn from populations subjected to long term variation in two key factors: socio-sexual and nutritional environments.

The social and sexual environment has emerged as an important driver of plasticity. For example, male Mediterranean fruit flies that perceive elevated sexual competition due to the presence of a conspecific male in the mating arena, transfer significantly more sperm to females during mating [21]. Similarly, male *D. melanogaster* respond to elevated sexual competition by mating for longer, transferring more of key seminal fluid proteins [22] and sperm [23-25] - thus achieving higher reproductive success [26]. Females also show related plastic responses to the presence of conspecifics [27, 28]. For example, in *D. melanogaster*, mated females are more aggressive towards rival females than are virgins [29, 30] and this aggression evolves to become stronger in populations maintained over time under strong female bias [30]. Female fecundity is also surprisingly plastic, and varies according to the pre- and post-mating social environment [31] [27]. For example, females maintained in same sex conspecific groups prior to mating lay significantly fewer eggs after mating than those kept alone, with the detection of deposits left by other females being key to the assessment of the female’s social environment [31]. These findings show that both sexes can respond plastically according to the level of potential competition experienced.

Plasticity in responses to sexual competition can also be strongly determined by an individual’s condition, which can vary with factors such as temperature, nutrition and population density. For example, in the moth *Plodia interpunctella*, males raised under high population density take longer to mature but have larger testes [32]. In *D. melanogaster*, increased availability of nutrition under lower larval density results in the production of larger females that are more aggressive [33]. However, once mated, it is smaller females that typically win physical contests [33]. In the Yellow dung fly (*Scathophaga stercoraria*) an increase in temperature is strongly correlated with a reduction in testes size [34]. These examples illustrate the influence of ‘conditional’ traits on the outcomes of sexual selection and competition [35]. Therefore, we predict that examination of the simultaneous effects of long term manipulations of condition and the social and sexual environment, as we <u>u</u>ndertook here, should be particularly informative for identifying the primary drivers of plasticity evolution.

Manipulation of the social and sexual environment can be achieved using phenotypic engineering of the immediate number, density or sex of potential competitors, of the sensory cues available [36, 37] or longer-term manipulations of the adult sex ratio (ASR, total number of adults in a population) over evolutionary time. ASR is a key variable to manipulate in the study of responses to the same and opposite sex competitive environment [38-40]. Sexual selection is predicted to become more intense in populations in which the ASR becomes unbalanced [38-42]. Under a male biased ASR, competition between females should decrease and between males should increase as males seek to maximise their reproductive success in the light of reduced mating opportunities [42, 43]. Consistent with this, experiments on *Tribolium castaneum* flour beetles have shown that males under intense competition maintained in male-biased regimes evolved more competitive sperm and gained higher paternity share in comparison to those from female-biased regimes [44]. Similarly, female biased ASRs are expected to have opposing effects, potentially increasing competition among females for access to males or resouces, while decreasing male-male sexual competition [45]. In line with this, in South African dung beetles of the genus *Onthophagus*, species without horns are typically female biased and occur at higher densities [46]. This is thought to have arisen because selection is focused on females fecundity rather than costly horn expression in males [47].

Recent work has extended evolutionary studies of the phenotypic and genetic responses to the effects of experimental variation in ASR and condition to include plasticity. This work shows that long term manipulations of ASR can provide sufficiently strong selection for plasticity to evolve. For example, studies on the lines used here showed that male courtship repertoire evolved in response to elevated male-male competition [48]. Here we used these same lines to test how the plastic fecundity responses of females to conspecifics evolve in response to the evolutionary manipulation of sexual selection and resource levels in adults. We asked whether the plastic responses in female fecundity [31] had evolved in experimental evolution regimes selected under male-biased (MB), equal sex (ES) and female-biased (FB) adult sex ratios maintained under high or low quality adult diets. The specific predictions were (i) that a history of elevated competition among females (in FB regimes) would select for increasingly sensitive plastic fecundity responses to the presence of conspecifics, which would (ii) be magnified under poor nutritional resource regimes.

## Methods

### Base stock maintenance and collection

All wild type non-focal flies were reared from wild-type Dahomey stock maintained at 25°C in a humidified room with a 12 h light: 12 h dark cycle. Flies were reared on a sugar-yeast-agar (SYA) medium (100g brewer’s yeast, 50g sucrose, 15g agar, 30mL Nipagin (10% solution), 3mL propionic acid, 0.97L water). Flies for use in experiments were grown by allowing females to first oviposit for 24 h on agar-grape juice plate (50g agar, 600mL red grape juice, 42mL Nipagin (10% solution), 1.1L water) to acclimatise, and then on a fresh agar-grape juice plate for 4 h. Larvae were collected from the 4 h egg collection plates and reared under a controlled density of 100 larvae per vial (24 x 75 mm) each containing 7ml SYA medium. At eclosion, adults were separated by sex within 6 h of eclosion to ensure virginity and stored 10 per vial in single sex groups in vials on standard SYA medium.

### Experimental evolution line maintenance

Females from the experimental evolution lines were derived from those used in previous studies [35, 43, 48, 49]. These lines comprise 3 independent replicates each of of EQual sex (EQ (50M: 50F)), Female Biased (FB (25M: 75F)) and 3 x Male biased (MB (70M: 30F)) lines, maintained as adults on either high or low SYA diets (100% versus 20% of the standard amount of yeast) (3 sex ratio regimes x 2 nutritional regimes x 3 replicates = 18 populations). Larvae from all regimes were always reared on standard the SYA medium each generation. MB lines were maintained under a slightly less extreme adult sex ratio than were FB lines, to ensure sufficient females to easily maintain the populations. Regimes were maintained as adults within plastic boxes (12cmW x 18cmL x 8.5cmD, with gauze lid) at 25°C in a humidified room with a 12 h light: 12 h dark cycle. Adults in the high yeast lines were given access to two fresh, standard SYA medium every two or three days, whilst the low yeast lines were similarly supplied with 20% SYA medium. Nine days after setting up the adults in the boxes, each line was supplied with an agar-grape juice egg collection plate, which was replaced on day 10. Egg collection plates were maintained at 25°C following their removal from the boxes and kept within in cotton bags for 2 days. 400 larvae were then picked from the egg collection plates and placed 100 per vial for each line. After eclosion of the adults from these vials, flies were anaesthetised using CO_2_, counted into the appropriate sex ratios, and placed again in plastic boxes. The lines have been maintained under these conditions since 23/12/2013. The flies used in the experiments were derived from generation 102 for block 1,& generation 109 for blocks 2 & 3.

### Collection of experimental females from the sex ratio lines

To minimise parental effects, and allow the detection of evolved responses, flies from the sex ratio lines for use in the experiments were reared under two generations of common garden conditions (equal sex ratio and nutritional conditions) for two generations prior to the set up of the experiments. To initiate these cultures, excess flies from the standard maintenance of the lines were transferred to 70ml bottles of standard SYA for 24 h, the adults then removed, and the deposited eggs allowed to mature to adulthood. Upon emergence the populations were transferred to an egg laying chamber (12cm diameter x 18cm high) and provided with an agar-grape juice egg collection plate for 24 h to acclimatise, which was then replaced with a fresh agar-grape juice egg collection plate for 4 h. Larvae were then picked at standard densities of 100 per vial into standard SYA vials. Upon eclosion adults were separated by sex to ensure virginity and stored 10 per vial on standard SYA medium.

### Effect of evolutionary manipulation of adult sex ratio and nutrition on socially plastic fecundity responses in females

Following the common garden rearing as described above, virgin females from all the lines were exposed to two social treatments, and maintained either alone or grouped with non-focal conspecific standard wild type females for 3 days and then mated. The experiment was carried out in two blocks (replicate population 1 first and replicate populations 2 & 3 simultaneously). All flies used in the experiment were aged between 8 -12 days old (from eclosion) at the time of mating. The grouped social treatments comprised 3 non-focal wild type virgin females + 1 focal sex ratio line virgin female, and the alone of 1 focal sex ratio virgin female held in social isolation. Non-focal wild type females for the grouped treatments were made identifyable by wing clipping under CO_2_ anaesthesia the day before introducing them to focal females. Females were held under these social treatments for 3 days until the mating assay, with 30 focal flies for each treatment combination.

Standard wild type males for the mating assay were transferred into fresh vials 24 h before the mating tests began. The mating assay set ups started at 9:00am and all matings were typically completed by 10:30am. Experimental females were each transferred into vials containing the wild type males for the mating assay via aspiration. Non-focal females were discarded, and the vacated vials were retained for subsequent counting of virgin eggs. Time of entry, mating latency and duration were all recorded, with any mating lasting less than 5min discarded, as they were likely to have been incomplete, with no sperm transfer [50]. After mating, males were removed and discarded. Focal females were retained in their mating vials for 24 h before being removed, eggs were then counted and kept 25°C in a humidified room with a 12 h light: 12 h dark cycle for 12 days. After 12 days all the emerged progeny were frozen and counted.

### Statistical analysis

All statistical analysis was performed using R-4.0.2 (R Core Team, 2020). All three replicates were analysed simultaneously, with the replicates (‘Population’) designated as a random factors. The Shapiro-Wilk test was used to check data were normally distributed and the Levene’s test to check the homogeneity of variances across treatments. Where data were not normally distributed, they were log_10_ transformed. Analysis of egg number, progeny number, latency and mating duration were performed using linear mixed effects models using the lme4 package [51]. Non-significant terms were dropped from the analysis by comparing models using Chi squared tests. The Akaike’s Information Criterion (AIC) was used to check for model fit. Post-hoc analysis was performed using the Tukey’s Test. Mating latency and duration were log_10_ transformed and analysed using linear mixed effects models. Linear models were conducted on the virgin egg counts. These data were initially analysed using the whole dataset. In subsequent analyses, zero egg counts (egg retaining females) were removed and the data for the egg laying females, as well as the number of egg retaining females were analysed separately, using a binomial generalized linear mixed model. For virgin egg data, we could not distinguish which eggs were laid by focal versus non focal females in the grouped treatments. Therefore, we analysed the data for alone versus grouped treatments separately, to avoid having to divide all of the grouped data replicates by 4 and thus compressing the variance of those data in comparison to the alone treatment.

## Results

### Evolution of plasticity in females is robust to evolutionary manipulations of adult sex ratio and nutrition

The main finding was that females from the sex ratio regimes retained plastic fecundity responses across all sex ratio and nutritional evolutionary treatments . However, counter to our predictions, this effect was not more marked in FB regimes and was also not magnified under poor resource conditions. In fact the extent of fecundity plasticity did not vary in magnitude across any treatments. These results are presented in full, below.

#### (i) Mating latency, duration, post-mating fecundity and egg to adult viability

Latency to mate was significantly longer overall in grouped compared to alone females (t = 2.418, residual DF = 877.72, *p=0*.*0158*; Fig. 1A; Fig. S1A-C). However, mating latency was not significantly different across evolutionary sex ratio or diet regimes. There was also no significant difference in mating duration of females held alone or in groups prior to mating, aross any of the sex ratio or diet regimes (t = -0.550, residual DF = 878.054 *p = 0*.*582*, Fig. 1B; Fig. S2A-C).

**Fig. 1.**
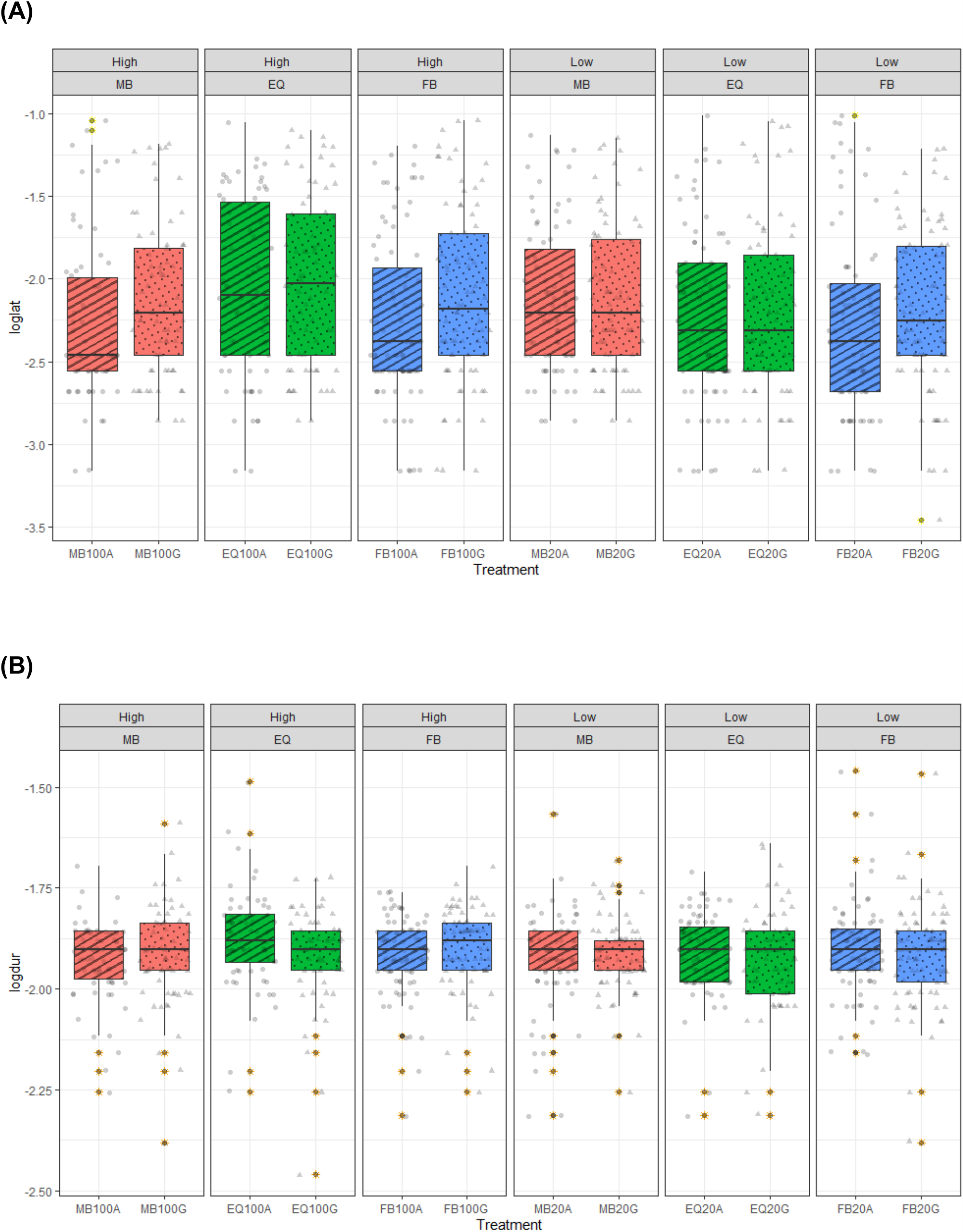
(A) Latency to mate (‘loglat’ in log_10_ minutes), and (B) mating duration (‘logdur’ in log_10_ minutes) of females from the sex ratio and diet regimes exposed for 3 days prior to mating to conspecifics or left alone. Females from the Female-biased (FB), Equal sex (EQ) or Male-biased (MB) sex ratios and standard 100% protein (High) or 20% protein (Low) diet regimes were tested. Females were either housed alone (A) or grouped with three rivals (G) for 3 days prior to mating. All conspecific non focal females and males were standard wild type. Boxplots show median line, with boxes representing upper and lower 25% quartile and whiskers representing the range, and points representing individual records, outliers highlighted in yellow.

Fecundity plasticity was retained in all treatments as shown by the finding that the number of eggs laid in the 24 h period after mating was significantly affected by the pre-mating social environment (t = -2.728, residual DF = 871.1465, *p=0*.*00649*), with grouped treatment females consistently laying fewer eggs after mating in comparison to the females held alone prior to mating (Fig. 2A; Fig. S3A-C) as has previously been reported in wild type females [31]. However, the magnitude of the plasticity in the fecundity responses of females to their pre-mating social environment was not significantly different across any of the sex ratio or diet regimes (table S1). Grouped females also produced fewer offspring than those held alone in the 24h after mating (t = -2.274, residual DF = 870.232, *p=0*.*0232*), again with no significant difference between sex ratio or diet regimes (Fig. 2B; Fig. S4A-C). There was no effect of social treatment, sex ratio or diet regime on egg-adult viability (percentage of post-mating eggs developing to adulthood after 12days) (Fig. 2C; Fig. S5A-C; table S2).

**Fig. 2.**
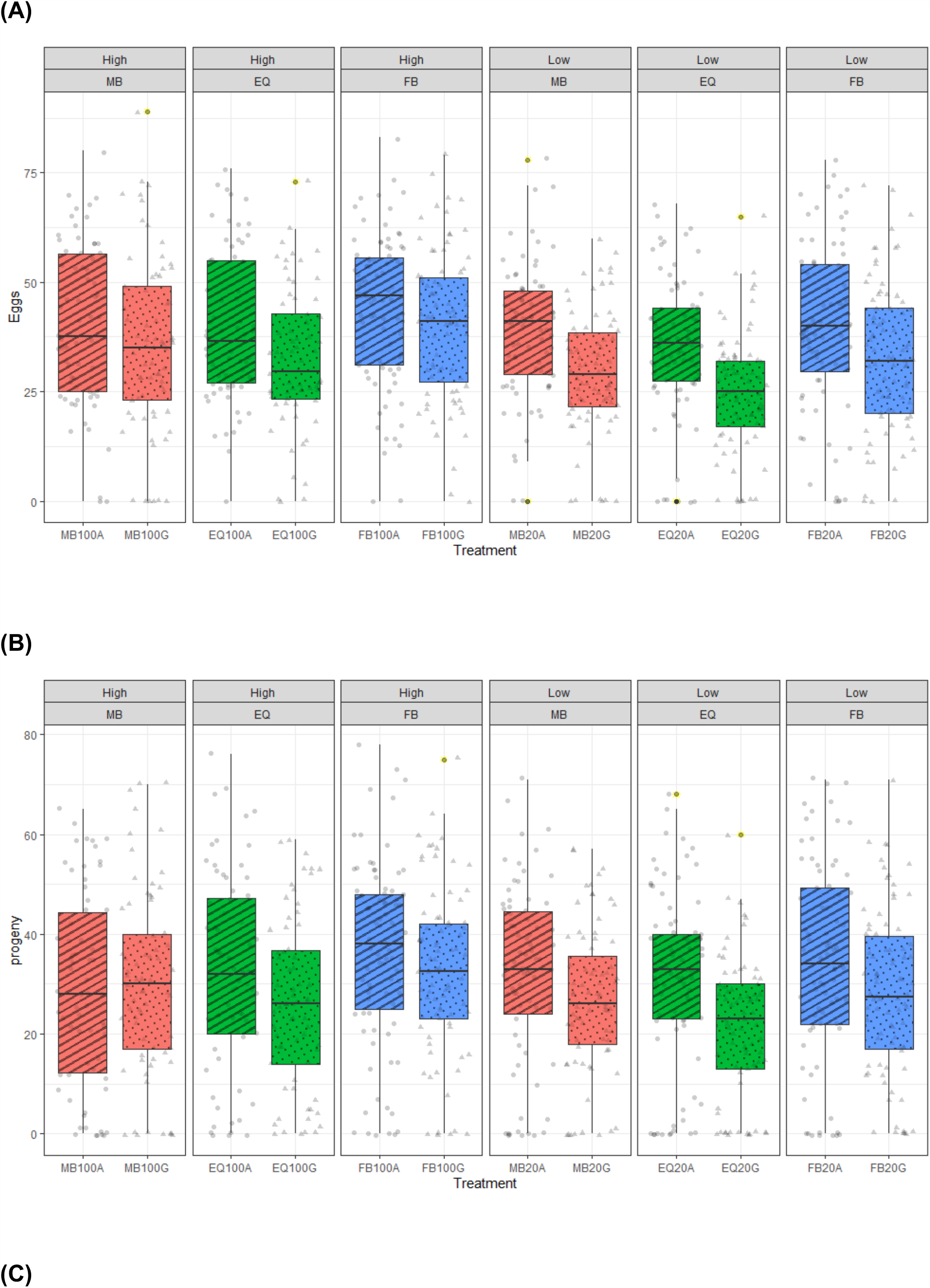
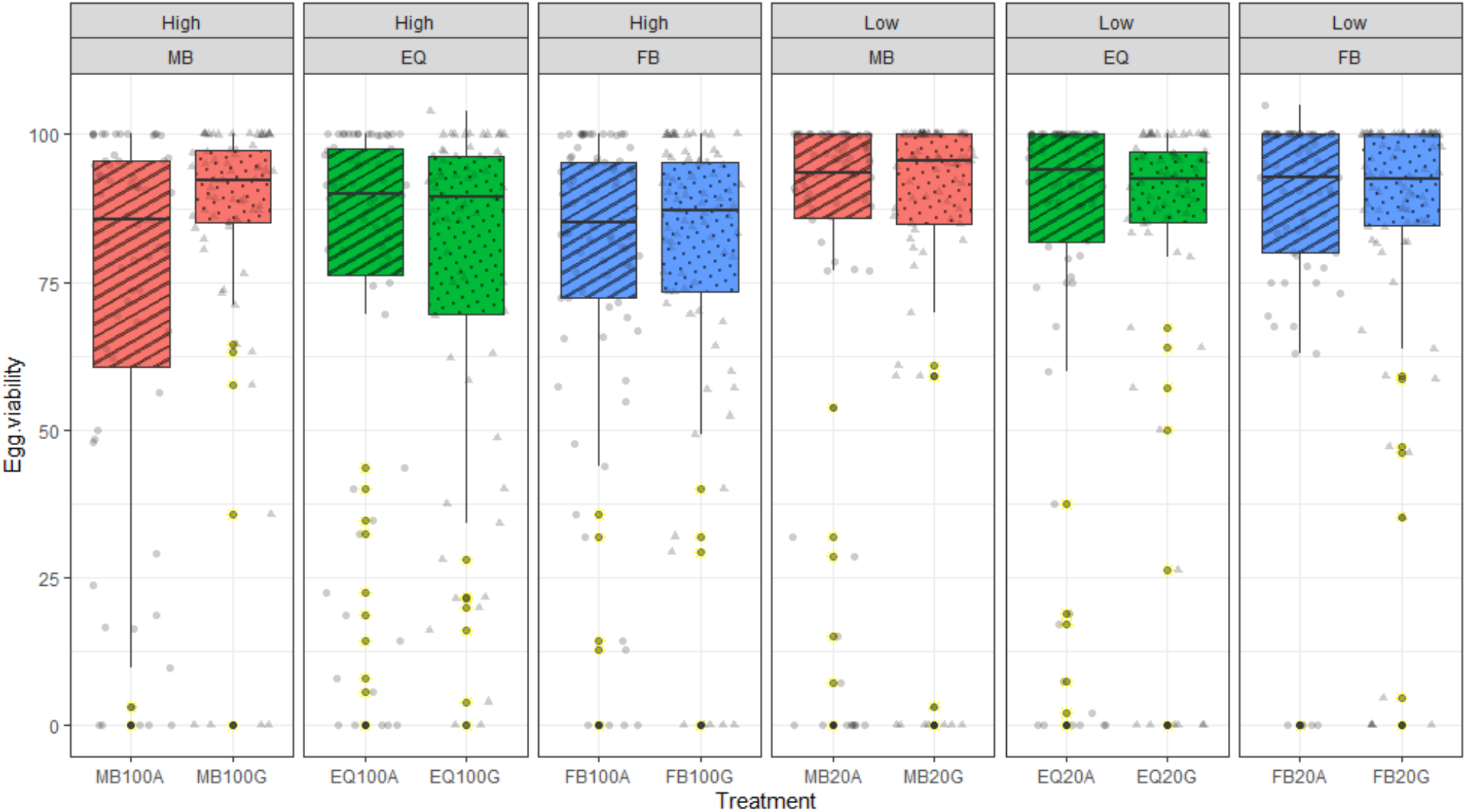
(A) Post-mating fecundity (‘Eggs’, number of eggs per female per 24h following mating), (B) Progeny production (‘progeny’, number of progeny emerging from the eggs laid in (A) per female per 24h following mating), (C) Egg to adult viability (‘Egg viability’, number of progeny / eggs per female per 24h following mating) of females from the sex ratio and diet regimes exposed for 3 days prior to mating to conspecifics or left alone. Females from the Female-biased (FB), Equal sex (EQ) or Male –biased (MB) sex ratios and standard protein (High) or 20% protein (Low) diet regimes were tested. Experimental females were either housed alone (A) or grouped with three rivals (G) prior to mating assay. All conspecific non focal females and males were standard wild type. Boxplots as per Fig.1.

The results show that fecundity plasticity was retained in all of the sex ratio regimes across both nutritional environments. However, counter to the prediction, the plasticity was not increased in females from FB regimes or magnified in females from the poor diet regimes.

#### (ii) Pre-mating virgin egg laying and egg retention

We also analysed the number of eggs laid, and number of eggs retained, by virgin females prior to mating in the alone versus grouped treatments. Separate analyses for the alone and grouped treatment data were conducted, as described above. Virgin egg counts for the alone treatment females differed significantly between food but not sex ratio regimes, with low food regime females held in social isolation prior to mating laying significantly higher numbers of virgin eggs than for high food regimes (t = 2.319, p = 0.0204; Fig. 3A; Fig. S6 A-C). In contrast, in the grouped treatment females there was no significant effect of sex ratio or dietary regime on the number of virgin eggs laid prior to mating (Fig. 3 B; Fig. S6D-F), though note that here we cannot distinguish those virgin eggs laid by the focal versus non focal females, which could have obscured variation among females. We separately analysed the frequency of vials with zero egg counts, to give an index of egg retention, though it is also possible that these were instances in which female ovaries contained no eggs. Among the grouped treatment females, there was again no significant difference in egg retention between females from the different sex ratios or dietary regimes. In the alone treatment females MB females from the low food regimes were significantly less likely to retain eggs in comparison to their high food counterparts (t = -1.538, p = 0.0004; Fig.4 A & B) an effect that was not observed for the other sex ratio regimes.

**Fig. 3.**
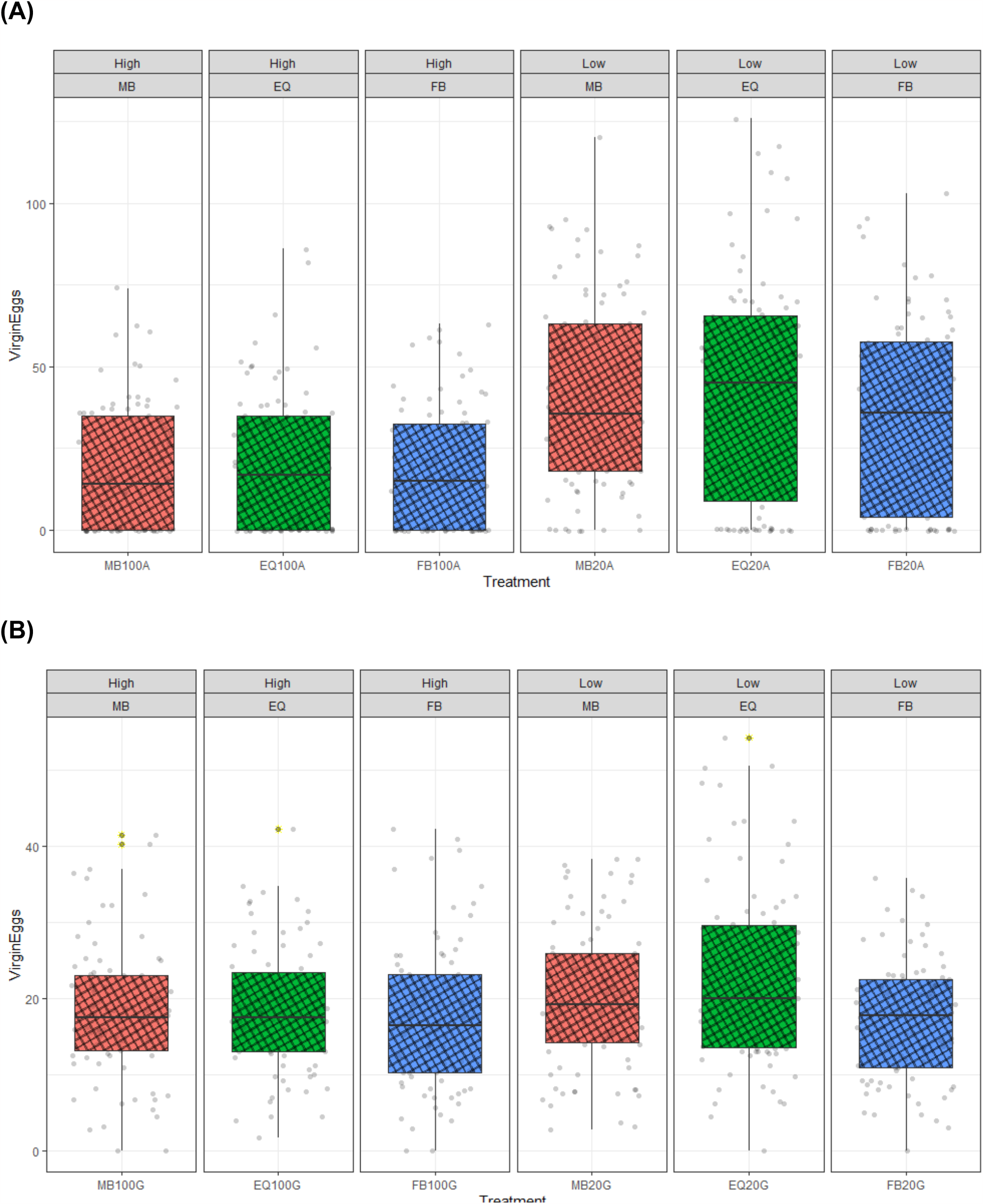
Virgin female fecundity for females from the sex ratio and diet lines held (A) alone and (B) in Groups. Virgin egg counts over 3 days prior to mating for females from the sex ratio and diet lines exposed for 3 days to conspecifics or left alone. Equal (EQ), Female-biased (FB), or Male–biased (MB) sex ratios from standard protein (High) or 20% protein (Low) diet regimes. Females were either housed alone (A) or grouped with three rivals (G). Boxplots showing median line, with boxes representing upper and lower 25% quartile and whiskers representing the range, and points representing individual records, outliers highlighted in yellow.

**Fig. 4.**
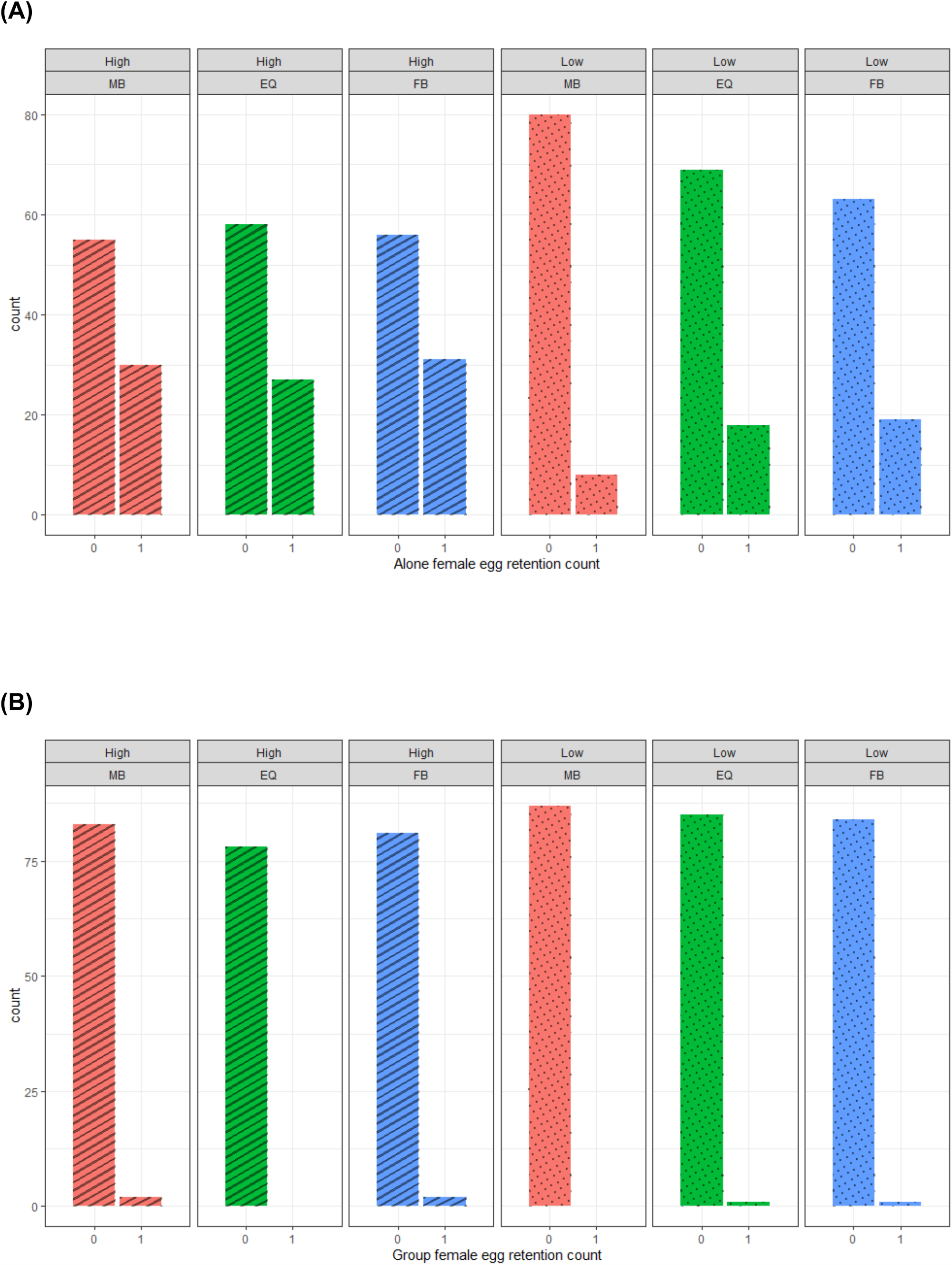
Number of females from the sex ratio lines retaining virgin eggs held (A) alone and (B) in groups. Females were scored as either “Egg layers” (egg retention count 0) who did produce virgin eggs over a 72h period prior to mating or as “egg retainers” (egg retention count 1), in which the female was still alive but did not produce any virgin eggs during this time. Focal females were exposed for 3 days to conspecifics (G) or left alone (A). Equal (EQ), Female-biased (FB), or Male –biased (MB) sex ratios from standard protein (High) or 20% protein (Low) diet regime. All conspecific non focal females were standard wild type.

Overall, these results showed significant responses of virgin egg laying to nutritional regime and of virgin egg retention to the adult sex ratio regime, but only in females held alone prior to mating.

## Discussion

Our main aim was to investigate whether plastic fecundity responses to the presence of conspecific females had evolved in lines with an evolutionary history of variation in sexual selection and adult diet availability. To test this idea, we compared the mating behaviour and fecundity plasticity of females drawn from MB, EQ and FB sex ratio environments maintained on high and low adult diets, raised through 2 generations of common garden rearing, and then housed alone or with three conspecific females prior to mating. Female plastic fecundity responses were retained in all regimes. However, neither plastic fecundity responses nor any pre-mating traits tested evolved in response to long term variation in the adult sex ratio or adult diet regimes. The results also revealed consistent plasticity in mating latency, with females from all regimes mating sooner when socially isolated prior to mating. The lack of differences in plasticity across the sex ratio or nutritional regimes regimes is not consistent with a lack of selection pressure, as we did see, among females socially isolated prior to mating, significant evolutionary responses of virgin egg laying to adult diet regime and of virgin egg retention to sex ratio regime.

### Mating latency, duration and fecundity plasticity

Though not part of the initial hypothesis we were testing, one outcome of the study was that mating latency was consistently plastic – females from all regimes mated significantly faster when they were held in isolation prior to mating. As neither adult diet nor sex ratio regime had any effect on this plasticity, it suggests that the response of female mating latency to the presence of conspecifics is robust across long term perturbations of the social and nutritional environment. The majority of females mated, which also suggests low variation among virgin females about whether to mate or not mate overall and that the sexual receptivity state of females to the cues signalled by males was not altered during the experimental evolution.

A consistent effect of mating latency according to the social environment is a new finding. It is possible that the general culturing procedures used to maintain the sex ratio and nutritional regimes used here conferred consistent benefits of plasticity in mating latency. This could be due to greater predictability of the conditions experienced by the experimental evolution regimes (specified densities, timings of culturing stages and non overlapping generations) in comparison to normal cage culture used for the wild type flies used in previous studies, in which this effect was not observed [31, 52]. We do not yet know the drivers of plasticity in mating latency. However, it is possible that females maintained on their own for three days prior to mating may perceive mating opportunities and competition for egg laying sites as low. This could increase a female’s willingness to mate rapidly with the first prospective partner encountered. In natural settings, *D. melanogaster* females tyically carry the sperm of at least two, but up to four different males [53]. Hence, if the opportunity to mate is perceived to be low, high receptivity to a first mating (i.e. quicker mating latency) could be more beneficial than waiting for any subsequent, even potentially fitter, male. We observed no differences in mating duration in females held alone or in groups prior to mating across any of the sexual selection or nutrition regimes. Hence, although females have the potential to influence mating duration [54], the results are consistent with previous reports that plasticity in extended mating duration is primarily under male control [26, 52, 55].

We set out to test whether fecundity plasticity evolved following long term variation in adult sex ratio and nutritional regimes. The results showed that this plasticity was retained across all evolutionary regimes, with females housed alone prior to mating producing significantly more eggs after mating than did females kept in groups. This is consistent with previous research [31]. Egg to adult viability did not differ across any regimes and hence, owing to their higher fecundity, females that were socially isolated prior to mating also produced significantly more offspring after mating than did those held in groups. Therefore, the differences observed in egg laying were wholly attributable to manipulations of the same sex social environment prior to mating, and not to the evolutionary lineage from which the focal females were drawn. Hence, conter to the main prediction, the plastic fecundity responses of females when exposed to conspecifics did not evolve according to variation in sexual selection or resource levels.

The benefits of fecundity plasticity are not yet clear. However, we suggest females held in groups prior to mating might perceive higher levels of resource competition, and thus lay fewer eggs after mating to reduce it. Following this reasoning, we had expected plasticity to evolve in response to elevated variation in sexual selection across the regimes [56]. Speccifically, we expected the increased level of female-female competition in FB regimes leading to enhanced plasticity in response to the same sex environment. The lack of response to sex ratio regime in reproductive output was surprising, given reported plasticity effects in males [26, 48, 57] and in other traits in females [49, 58-60]. The results are consistent with the idea that the maintenance of this type of plasticity may carry low fitness costs, or that any such cost is accumulated across lifespan [58, 60-63], which would not have been captured here.

Fecundity plasticity was also expected to respond differentially to the nutritional regimes. Our hypothesis was that it would be costly for females to oviposit eggs in an area in which other females were doing the same, particularly under food limitation. Under these conditions, females should retain eggs or search for less densely populated oviposition sites. This was not observed, as the extent of fecundity plasticity was unaffected by nutritional regimes. Given that protein restrictive diets can also reduce protein content in eggs [64] and that low protein in adult diets generally significantly reduces female fecundity [65, 66], we expected that the lines maintained under an evolutionary history of restrictive adult diets (20% protein) would evolve to produce fewer eggs or alternatively to become more efficient in nutrient acquisition leading to higher fecundity when measured in a common garden. There was little evidence that this was the case, with no significant differences in post-mating fecundity attributable to evolution under the different adult dietary regimes. This may indicate that the females are either gaining sufficient protein from the carry over effects of resources gained during larval development [67], at least for an initial batch of eggs [67, 68] or that the restricted adult diet lines have evolved to cope with limited protein availability in adulthood without reducing egg quality [64].

### Evolutionary responses of virgin egg laying and egg retention

The lack of predicted responses in fecundity plasticity could be explained by lack of sufficient selection pressure. However, arguing against this were the evolutionary responses to nutritional and sex ratio regimes that we did observe in virgin egg laying and egg retention. Interestingly, these responses were evident only among females that were socially isolated prior to mating.

The number of virgin eggs laid by focal females kept alone was significantly higher for females drawn from the low adult food regimes, but was unaffected by adult sex ratio regime. This effect could result from elevated selection for food utilisation efficiency [68] among females exposed over the long term to a poor adult nutritional environment. Why it would be evident only in females held in isolation prior to mating is not clear. However, we note that it was not possible to designate the virgin eggs of the focal females when they were held in groups, which could have obscured any variation in egg laying among focal females. Females held in groups also have the potential to learn from each other regarding oviposition decisions [28], which would be interesting to test further.

Among females held in isolation prior to mating, there was also a significant effect of the sex ratio regime on virgin egg retention, with MS females from the low food regimes being more likely to retain their eggs, or to have no eggs to lay, in comparison to MB females from the high food regimes. No such differences was seen for females from the other sex ratio regimes. The result showed that virgin egg retention was contingent on both the long term sex ratio and nutritional regime. The response of increased egg retention in MB females on low in comparison to high food could indicate that virgin egg production for these females in particular is potentially costly. However, the significance of virgin egg production overall is not well understood and copsts of virgin egg production are generally assumed to be low [69].

Overall, consistent with previous works, we show that females adjust their post-mating fecundity according to the social environment.This plasticity was unexpectedly robust to long term evolutionary manipulations of sexual selection and resource levels. The results show how such plasticity can to be fairly hard wired to evolutionary perturbations.

## Acknowledgements

We thank Emily K Fowler, Alice Dore, Stewart Leigh, Wayne G Rostant for help with mating assays and stimulating conversations regarding the manuscript.

## Author’s contributions

T.C., W.H., M.J.G.G and N.M conceptualised the study. N. M. conducted the labwork and the data analyses and wrote the first draft of the manuscript. All authors contributed to writing and revising the manuscript and approved the final version.

## Data availability

DRYAD link to raw data (doi to be appended). Supplementary figures and tables are presented in Additional file 1.

## Declarations

### Ethics approval and consent to participate

Protocols were approved by the appropriate animal welfare and ethical review board of the university. Formal ethical approval was not required. All methods were incompliance with relevant guidelines.

### Consent for publication

Not applicable

## References

1. Bretman A, Gage MJ, Chapman T. Quick-change artists: male plastic behavioural responses to rivals. Trends in Ecol Evol. 2011;26:467–73.

2. Wedell N, Gage MJG, Parker GA. Sperm competition, male prudence and sperm-limited females. Trends in Ecology & Evolution. 2002;17(7):313–20. PubMed PMID: ISI:000176331800011.

3. Kasumovic MM, Brooks RC. It’s all who you know: the evolution of socially cued anticipatory plasticity as a mating strategy. Q Rev Biol. 2011;86(3):181–97. doi: 10.1086/661119. PubMed PMID: 21954701.

4. Sheehy KA, Laskowski KL. Correlated behavioural plasticities: insights from plasticity evolution, the integrated phenotype and behavioural syndromes. Animal Behaviour. 2023;10.1016/j.anbehav.2023.04.007.

5. Dingemanse NJ, Wolf M. Between-individual differences in behavioural plasticity within populations: causes and consequences. Animal Behaviour. 2013;85:1031–9.

6. Snell-Rood EC. An overview of the evolutionary causes and consequences of behavioural plasticity. Animal Behaviour. 2013;85:1004–11.

7. Moczek AP, Sultan S, Foster S, Ledon-Rettig C, Dworkin I, Nijhout HF, et al. The role of developmental plasticity in evolutionary innovation. Proc Biol Sci. 2011;278(1719):2705–13. Epub 20110615. doi: 10.1098/rspb.2011.0971. PubMed PMID: 21676977; PubMed Central PMCID: PMCPMC3145196.

8. Pfennig DW. Phenotypic plasticity & evolution: causes, consequences, controversies: Taylor and Francis; 2021. 436 p.

9. Price TD, Qvarnstrom A, Irwin DE. The role of phenotypic plasticity in driving genetic evolution. Proc Biol Sci. 2003;270(1523):1433–40. doi: 10.1098/rspb.2003.2372. PubMed PMID: 12965006; PubMed Central PMCID: PMCPMC1691402.

10. Van Buskirk J. Behavioural plasticity and environmental change. In: Candolin U, Wong BBM, editors. Behavioural Responses to a Changing World Mechanisms and Consequences: Oxford University Press; 2012. p. 145–58.

11. Snell-Rood EC, Davidowitz G, Papaj DR. Plasticity in learning causes immediate and transgenerational changes in allocation of resources. Integr Comp Biol. 2013;53(2):329–39. Epub 20130426. doi: 10.1093/icb/ict030. PubMed PMID: 23624867.

12. Lange EC, Erk S, Ptacek MB, Travis J, Hughes KA. Socially cued anticipatory plasticity predicts male primary mating tactic but not mating behaviour rates. Animal Behaviour,. 2023;196:43–53.

13. Yoon KJ, Cunningham CB, Bretman A, Duncan EJ. One genome, multiple phenotypes: decoding the evolution and mechanisms of environmentally induced developmental plasticity in insects. Biochem Soc Trans. 2023;51(2):675–89. doi: 10.1042/BST20210995. PubMed PMID: 36929376.

14. Zajitschek F, Zajitschek S, Canton C, Georgolopoulo G, Friberg U, Maklakov AA. Evolution under dietary restriction increases male reproductive performance without survival cost. . Proc Roy Soc B. 2016;283:1825.

15. Zajitschek F, Zajitschek SR, Friberg U, Maklakov AA. Interactive effects of sex, social environment, dietary restriction, and methionine on survival and reproduction in fruit flies. Age. 2013;35(4):1193–204. Epub 20120715. doi: 10.1007/s11357-012-9445-3. PubMed PMID: 22798158; PubMed Central PMCID: PMCPMC3705097.

16. Bretman A, Chapman T, Rouse J, Wigby S. Playing to the crowd: Using Drosophila to dissect mechanisms underlying plastic male strategies in sperm competition games. Advances in the Study of Behavior. 2023;55:1–35.

17. Collins DH, Prince DC, Donelan JL, Chapman T, Bourke AFG. Developmental diet alters the fecundity-longevity relationship and age-related gene expression in Drosophila melanogaster. bioRxiv. 2023;10.1101/2023.01.16.524185

18. Bretman A, Westmancoat JD, Gage MJG, Chapman T. Individual plastic responses by males to rivals reveal mismatches between behaviour and fitness outcomes. Proc Roy Soc B,. 2012;279:2868–76.

19. Scheiner SM, Levis NA. The loss of phenotypic plasticity via natural selection: genetic assimilation. In: Pfennig DW, editor. Phenotypic Plasticity & Evolution: CRC Press; 2021. p. 161–81.

20. Lange EC, Travis J, Hughes KA, M’Gonigle LK. Can You Trust Who You See? The Evolution of Socially Cued Anticipatory Plasticity. Am Nat. 2021;197(4):E129–E42. Epub 20210211. doi: 10.1086/712919. PubMed PMID: 33755539.

21. Gage MJG. Risk of sperm competition directly affects ejaculate size in the Mediterranean fruit fly. Anim Behav. 1991;44:1036–7.

22. Wigby S, Sirot LK, Linklater JR, Buehner N, Calboli FCF, Bretman A, et al. Seminal fluid protein allocation and male reproductive success Current Biology. 2009;19:1–7.

23. Moatt JP, Dytham C, Thom MD. Sperm production responds to perceived sperm competition risk in male Drosophila melanogaster. Physiol Behav. 2014;131:111–4. Epub 20140424. doi: 10.1016/j.physbeh.2014.04.027. PubMed PMID: 24769021.

24. Hopkins BR, Sepil I, Thezenas ML, Craig JF, Miller T, Charles PD, et al. Divergent allocation of sperm and the seminal proteome along a competition gradient in Drosophila melanogaster. Proceedings of the National Academy of Sciences of the United States of America. 2019;116(36):17925–33. Epub 20190820. doi: 10.1073/pnas.1906149116. PubMed PMID: 31431535; PubMed Central PMCID: PMCPMC6731677.

25. Garbaczewska M, Billeter JC, Levine JD. Drosophila melanogaster males increase the number of sperm in their ejaculate when perceiving rival males. J Insect Physiol. 2013;59(3):306–10. Epub 20121120. doi: 10.1016/j.jinsphys.2012.08.016. PubMed PMID: 23178803.

26. Bretman A, Fricke C, Chapman T. Plastic responses of male D. melanogaster to the level of sperm competition increase male reproductive fitness. Proc Roy Soc B. 2009;276:1705–11.

27. Bailly TP, Kohlmeier P, Etienne RS, Wertheim B, Billeter JC. Social modulation of oogenesis and egglaying in Drosophila melanogaster. bioRxiv. 2021.

28. Sarin S, Dukas R. Social learning about egg-laying substrates in fruitflies. Proc Biol Sci. 2009;276(1677):4323–8. Epub 20090916. doi: 10.1098/rspb.2009.1294. PubMed PMID: 19759037; PubMed Central PMCID: PMCPMC2817106.

29. Nilsen SP, Chan YB, Huber R, Kravitz EA. Gender-selective patterns of aggressive behavior in Drosophila melanogaster. Proceedings of the National Academy of Sciences. 2004;101:12342–7.

30. Bath E, Bowden S, Peters C, Reddy A, Tobias JA, Easton-Calabria E, et al. Sperm and sex peptide stimulate aggression in female Drosophila. Nature Ecology and Evolution. 2017.

31. Fowler EK, Leigh S, Rostant WG, Thomas A, Bretman A, Chapman T. Memory of social experience affects female fecundity via perception of fly deposits. BMC Biol. 2022;20(1):244. Epub 20221031. doi: 10.1186/s12915-022-01438-5. PubMed PMID: 36310170; PubMed Central PMCID: PMCPMC9620669.

32. Gage MJG. Continuous variation in reproductive strategy as an adaptive reponse to population density in the moth Plodia interpunctella. Proc Roy Soc Lond B. 1995;261:25–30.

33. Bath E, Morimoto J, Wigby S. The developmental environment modulates mating-induced aggression and fighting success in adult female Drosophila. Funct Ecol. 2018;32(11):2542–52. Epub 20180928. doi: 10.1111/1365-2435.13214. PubMed PMID: 31007331; PubMed Central PMCID: PMCPMC6472669.

34. Bernasconi G, Hellriegel B, Heyland A, Ward PI. Sperm survival in the female reproductive tract in the fly Scathophaga stercoraria (L.). Journal of Insect Physiology. 2002;48(2):197–203. PubMed PMID: ISI:000174738100007.

35. Bath E, Edmunds D, Norman J, Atkins C, Harper L, Rostant WG, et al. Sex ratio and the evolution of aggression in fruit flies. Proc Biol Sci. 2021;288(1947):20203053. Epub 20210317. doi: 10.1098/rspb.2020.3053. PubMed PMID: 33726599; PubMed Central PMCID: PMCPMC8059548.

36. Bretman A, Fricke C, Hetherington P, Stone R, Chapman T. Variation in exposure to rivals and plastic responses to sperm competition in Drosophila melanogaster. Behav Ecol. 2010;21: 317–21.

37. Bretman A, Westmancoat JD, Gage MJG, Chapman T. Multiple, redundant cues used by males to detect mating rivals. Curr Biol. 2011;21:617–22.

38. Emlen ST, Oring LW. Ecology, sexual selection, and the evolution of mating systems. Science. 1977;197:215–23.

39. Clutton-Brock TH. Reproductive success : studies of individual variation in contrasting breeding systems. Chicago: University of Chicago Press; 1988.

40. Clutton-Brock TH. Sexual selection in males and females. Science. 2007;318:1882–5

41. Pitnick S. Operational sex-ratios and sperm limitation in populations of drosophila-pachea. Behavioral Ecology And Sociobiology 33. 1993:383–91.

42. Hollis B, Koppik M, Wensing KU, Ruhmann H, Genzoni E, Erkosar B, et al. Sexual conflict drives male manipulation of female postmating responses in Drosophila melanogaster. Proceedings of the National Academy of Sciences of the United States of America. 2019;116(17):8437–44. Epub 20190408. doi: 10.1073/pnas.1821386116. PubMed PMID: 30962372; PubMed Central PMCID: PMCPMC6486729.

43. Sepil I, Perry JC, Dore A, Chapman T, Wigby S. Experimental evolution under varying sex ratio and nutrient availability modulates male mating success in Drosophila melanogaster. Biol Lett. 2022;18(6):20210652. Epub 20220601. doi: 10.1098/rsbl.2021.0652. PubMed PMID: 35642384; PubMed Central PMCID: PMCPMC9156920.

44. Godwin JL, Vasudeva R, Michalczyk Ł, Martin OY, Lumley AJ, Chapman T, et al. Experimental evolution reveals that sperm competition intensity selects for longer, more costly sperm. Evolution Letters. 2017;1:102–13.

45. Holveck MJ, Gauthier AL, Nieberding CM. Dense, small and male-biased cages exacerbate male– male competition and reduce female choosiness in Bicyclus anynana. Animal Behaviour. 2015;104:229–45.

46. Pomfret JC, Knell RJ. Crowding, sex ratio and horn evolution in a South African beetle community. Proc Biol Sci. 2008;275(1632):315–21. doi: 10.1098/rspb.2007.1498. PubMed PMID: 18048281; PubMed Central PMCID: PMCPMC2593729.

47. Simmons LW, Kvarnemo C. Costs of breeding and their effects on the direction of sexual selection. Proc Biol Sci. 2006;273(1585):465–70. doi: 10.1098/rspb.2005.3309. PubMed PMID: 16615214; PubMed Central PMCID: PMCPMC1560200.

48. Dore AA, Rostant WG, Bretman A, Chapman T. Plastic male mating behaviour evolves in response to the competitive environment. Evolution. 2020;75:101–15

49. Rostant WG, Mason JS, deCoriolis J-C, Chapman T. Evolution of lifespan and ageing in response to sexual conflict is sex-specific and condition-dependent. Evolution Letters. 2020;4:54–64.

50. Gilchrist AS, Partridge L. Why it is difficult to model sperm displacement in Drosophila melanogaster: The relation between sperm transfer and copulation duration. EVOLUTION. 2000;54:534–42.

51. Bates DMBS, Mächler M, Bolker B, Walker S. Fitting linear mixed-effects models using lme4. J Stat Softw. 2015;2015:1–48.

52. Fowler EK, Leigh SA, Bretman A, Chapman T. Reproductive plasticity in both sexes interacts to determine mating behaviour and fecundity. Evolution. 2022;76:2116–29.

53. Imhof M, Harr B, Brem G, Schlotterer C. Multiple mating in wild Drosophila melanogaster revisited by microsatellite analysis. Molecular Ecology 7. 1998:915–7.

54. Lefranc A, Bundgaard J. The influence of male and female body size on copulation duration and fecundity in Drosophila melanogaster. Hereditas. 2000;132:243–7.

55. Bretman A, Westmancoat JD, Chapman T. Male control of mating duration following exposure to rivals in fruitflies. J Insect Physiol. 2013;59:824–7.

56. Rosvall KA. Intrasexual competition in females: evidence for sexual selection? Behav Ecol. 2011;22(6):1131–40. Epub 20110908. doi: 10.1093/beheco/arr106. PubMed PMID: 22479137; PubMed Central PMCID: PMCPMC3199163.

57. Dore AA, Bretman A, Chapman T. Fitness consequences of redundant cues of competition in male D. melanogaster. Ecology and Evolution. 2020;10.22541/au.158030902.29897139.

58. Holland B, Rice WR. Experimental removal of sexual selection reverses intersexual antagonistic coevolution and removes a reproductive load. Proceedings of the National Academy of Sciences of the United States of America. 1999;96:5083–8.

59. House CM, Rapkin J, Hunt J, Hosken DJ. Operational sex ratio and density predict the potential for sexual selection in the broad-horned beetle. Animal Behaviour. 2019;152:63–9.

60. Wigby S, Chapman T. Female resistance to male harm evolves in response to manipulation of sexual conflict. Evolution. 2004;58:1028–37.

61. Nandy B, Chakraborty P, Gupta V, Ali SZ, Prasad NG. Sperm competitive ability evolves in response to experimental alteration of operational sex ratio. Evolution. 2013;67(7):2133–41. Epub 20130321. doi: 10.1111/evo.12076. PubMed PMID: 23815666.

62. Tilszer M, Antoszczyk K, Salek N, Zajac E, Radwan J. Evolution under relaxed sexual conflict in the bulb mite Rhizoglyphus robini. Evolution. 2006;60(9):1868–73. doi: 10.1554/06-060.1. PubMed PMID: 17089971.

63. Chapman T, Liddle LF, Kalb JM, Wolfner MF, Partridge L. Cost of mating in. Drosophila melanogaster. females is mediated by male accessory gland products. Nature. 1995;373:241–4.

64. Kutzer MA, Armitage SA. The effect of diet and time after bacterial infection on fecundity, resistance, and tolerance in Drosophila melanogaster. Ecol Evol. 2016;6(13):4229–42. Epub 20160525. doi: 10.1002/ece3.2185. PubMed PMID: 27386071; PubMed Central PMCID: PMCPMC4884575.

65. Dick KB, Ross CR, Yampolsky LY. Genetic variation of dietary restriction and the effects of nutrient-free water and amino acid supplements on lifespan and fecundity of Drosophila. Genet Res (Camb). 2011;93(4):265–73. Epub 20110718. doi: 10.1017/S001667231100019X. PubMed PMID: 21767463.

66. Zajitschek F, Georgolopoulos G, Vourlou A, Ericsson M, Zajitschek SRK, Friberg U, et al. Evolution Under Dietary Restriction Decouples Survival From Fecundity in Drosophila melanogaster Females. J Gerontol A Biol Sci Med Sci. 2019;74(10):1542–8. doi: 10.1093/gerona/gly070. PubMed PMID: 29718269.

67. Aguila JR, Hoshizaki DK, Gibbs AG. Contribution of larval nutrition to adult reproduction in Drosophila melanogaster. J Exp Biol. 2013;216(Pt 3):399–406. Epub 20121004. doi: 10.1242/jeb.078311. PubMed PMID: 23038728.

68. Bowman E, Tatar M. Reproduction regulates Drosophila nutrient intake through independent effects of egg production and sex peptide: Implications for aging. Nutr Healthy Aging. 2016;4(1):55–61. Epub 20161027. doi: 10.3233/NHA-1613. PubMed PMID: 28035342; PubMed Central PMCID: PMCPMC5166518.

69. Tatar M, Promislow DEL. Fitness Costs of Female Reproduction. Evolution. 1997;51(4):1323–6. doi: 10.1111/j.1558-5646.1997.tb03980.x. PubMed PMID: 28565490.

