## Supplementary material for "Socially plastic responses in females are robust to evolutionary manipulations of adult sex ratio and adult nutrition": Supplementary information Social_BMC.pdf

Figures S1-S5

Tables S1-S4

A

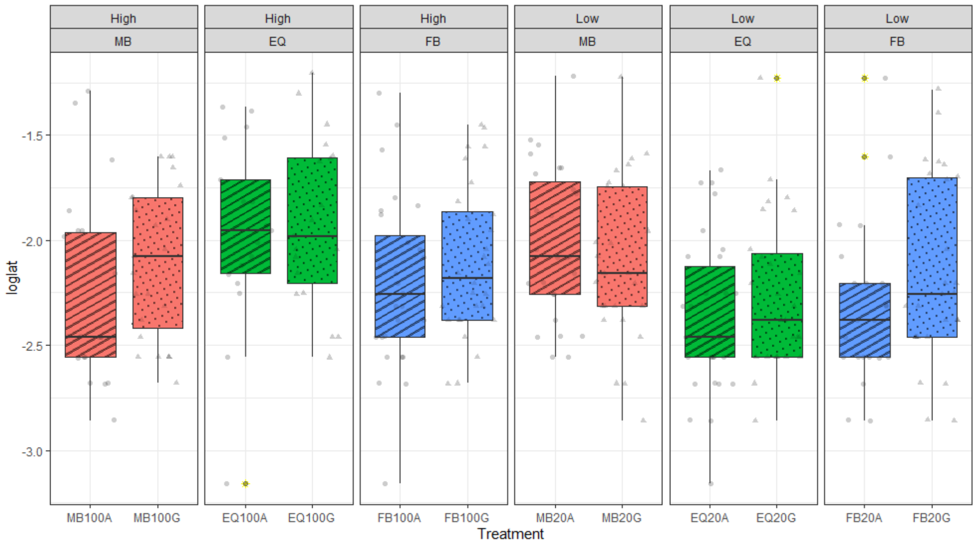
High
MB
High
EQ
High
FB
Low
MB
Low
EQ
Low
FB
-1.5
-2.0
-2.5
-3.0
loglat
MB100A
MB100G
EQ100A
EQ100G
FB100A
FB100G
MB20A
MB20G
EQ20A
EQ20G
FB20A
FB20G
Treatment

B

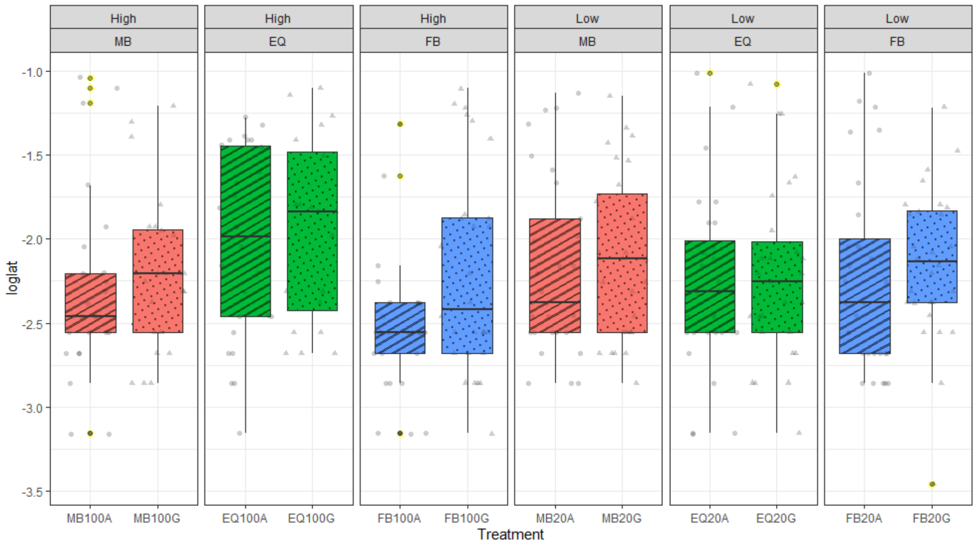
High
MB
High
EQ
High
FB
Low
MB
Low
EQ
Low
FB
-1.0
-1.5
-2.0
-2.5
-3.0
-3.5
loglat
MB100A
MB100G
EQ100A
EQ100G
FB100A
FB100G
MB20A
MB20G
EQ20A
EQ20G
FB20A
FB20G
Treatment

C

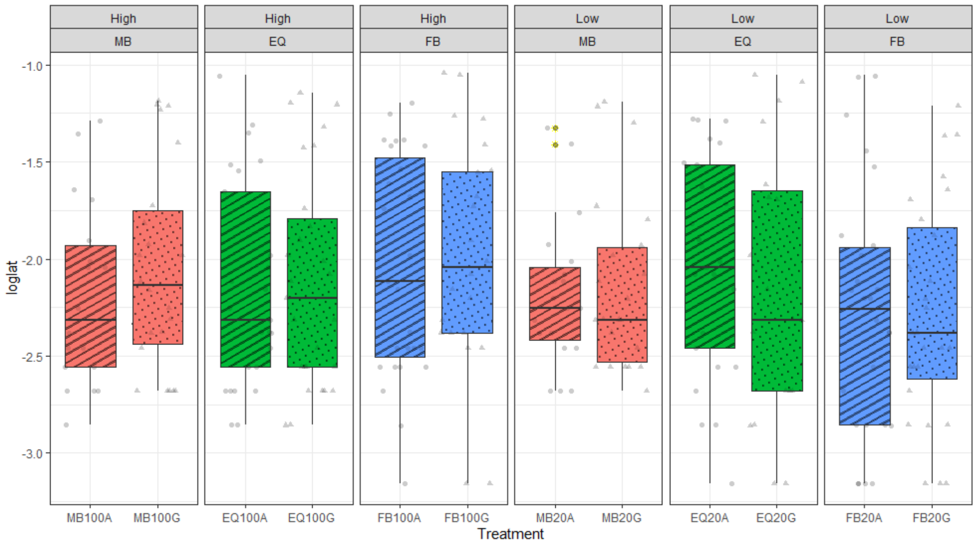
High
MB
High
EQ
High
FB
Low
MB
Low
EQ
Low
FB
-1.0
-1.5
-2.0
-2.5
-3.0
loglat
MB100A
MB100G
EQ100A
EQ100G
FB100A
FB100G
MB20A
MB20G
EQ20A
EQ20G
FB20A
FB20G
Treatment

**Figure S1, A, B, C Individual replicate population Latency to mate ($log_{10}$ minutes), of females from the sex ratio and diet regimes exposed for 3 days prior to mating to conspecifics or left alone.** Females from the Female-biased (FB), Equal sex (EQ) or Male –biased (MB) sex ratios and standard protein (High) or 20% protein (Low) diet regimes were tested. Females were either housed alone (A) or grouped with three rivals (G) for 3 days prior to mating. All conspecific non focal females and males were standard wild type. Boxplots show median line, with boxes representing upper and lower 25% quartile and whiskers representing the range, and points representing individual records, outliers highlighted in yellow.

A

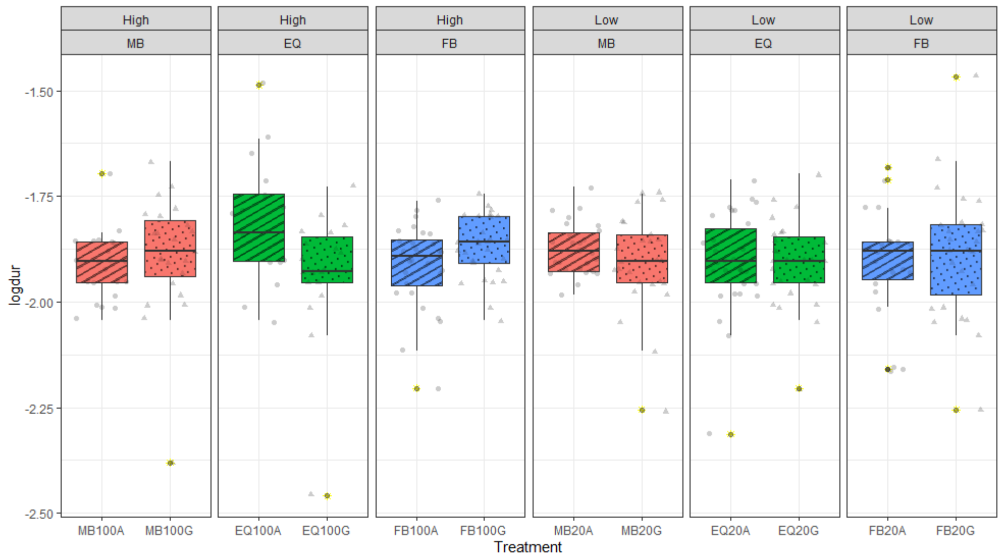
High
MB
High
EQ
High
FB
Low
MB
Low
EQ
Low
FB
-1.50
-1.75
-2.00
-2.25
-2.50
logdur
MB100A
MB100G
EQ100A
EQ100G
FB100A
FB100G
MB20A
MB20G
EQ20A
EQ20G
FB20A
FB20G
Treatment

B

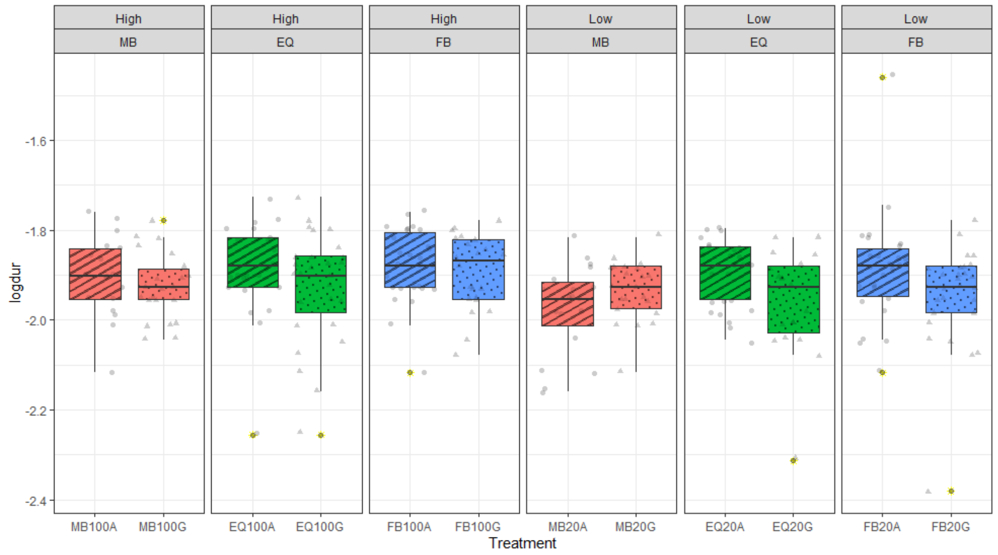
High
MB
High
EQ
High
FB
Low
MB
Low
EQ
Low
FB
-1.6
-1.8
-2.0
-2.2
-2.4
logdur
MB100A
MB100G
EQ100A
EQ100G
FB100A
FB100G
MB20A
MB20G
EQ20A
EQ20G
FB20A
FB20G
Treatment

C

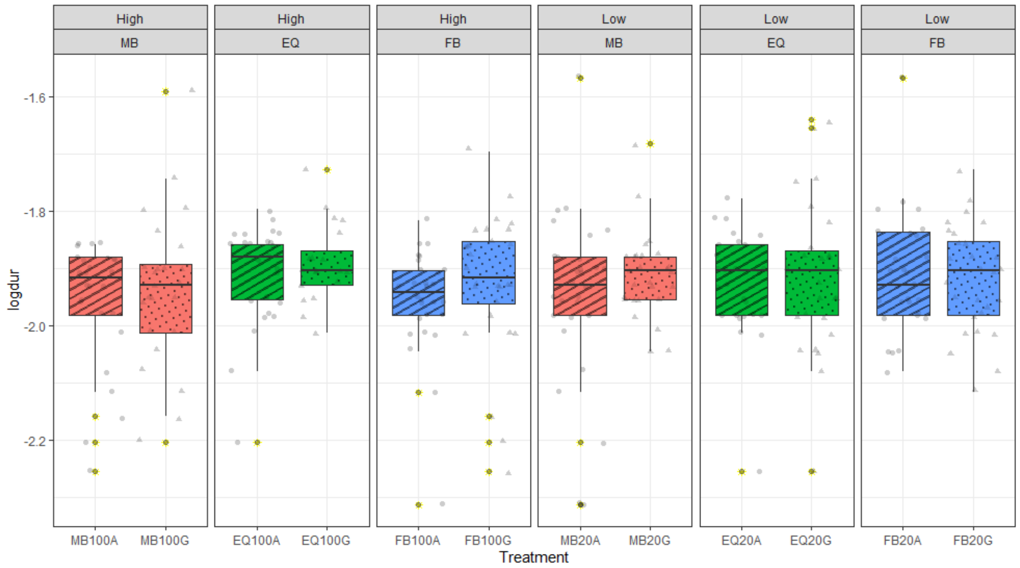
High
MB
High
EQ
High
FB
Low
MB
Low
EQ
Low
FB
-1.6
-1.8
-2.0
-2.2
logdur
MB100A
MB100G
EQ100A
EQ100G
FB100A
FB100G
MB20A
MB20G
EQ20A
EQ20G
FB20A
FB20G
Treatment

**Figure S2, A, B, C. Individual replicate population figures for mating duration ($log_{10}$ minutes), of females from the sex ratio and diet regimes exposed for 3 days prior to mating to conspecifics or left alone.** Females from the Female-biased (FB), Equal sex (EQ) or Male –biased (MB) sex ratios and standard protein (High) or 20% protein (Low) diet regimes were tested. Females were either housed alone (A) or grouped with three rivals (G) for 3 days prior to mating. All conspecific non focal females and males were standard wild type. Boxplots as per Fig.S1.

A

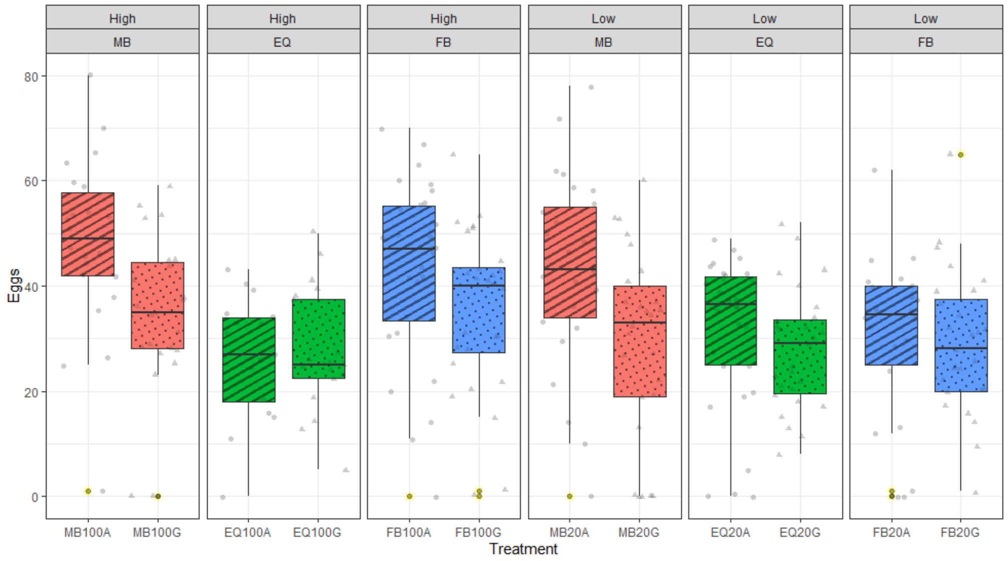
High
MB
High
EQ
High
FB
Low
MB
Low
EQ
Low
FB
Eggs
80
60
40
20
0
MB100A
MB100G
EQ100A
EQ100G
FB100A
FB100G
MB20A
MB20G
EQ20A
EQ20G
FB20A
FB20G
Treatment

B

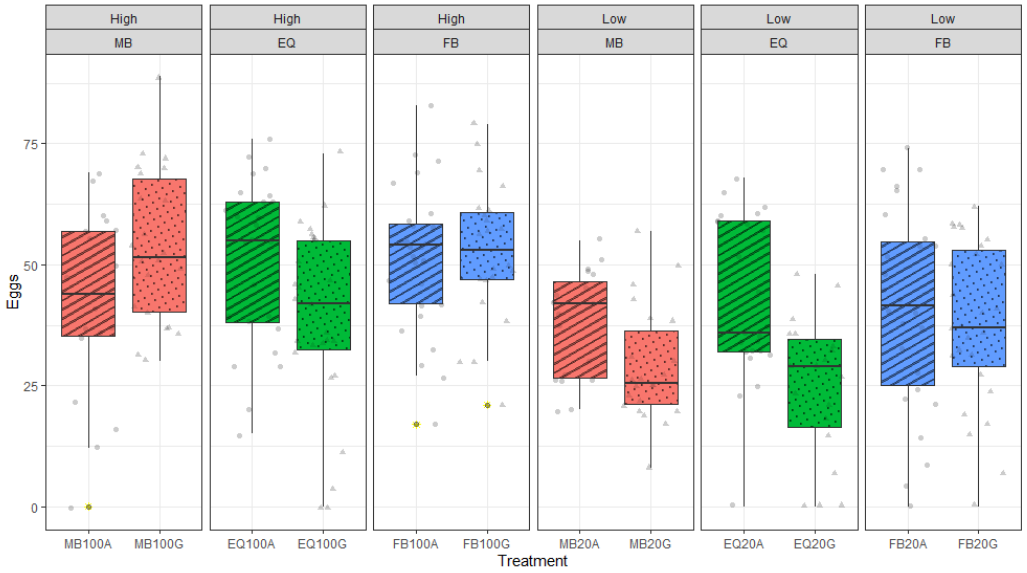
High
MB
High
EQ
High
FB
Low
MB
Low
EQ
Low
FB
Eggs
75
50
25
0
MB100A
MB100G
EQ100A
EQ100G
FB100A
FB100G
MB20A
MB20G
EQ20A
EQ20G
FB20A
FB20G
Treatment

C

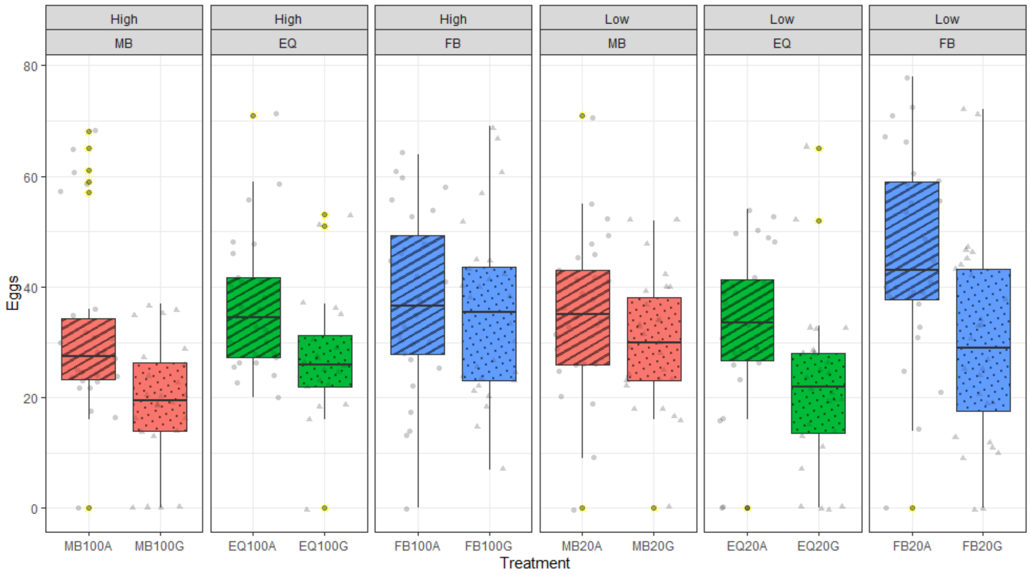
High
MB
High
EQ
High
FB
Low
MB
Low
EQ
Low
FB
Eggs
80
60
40
20
0
MB100A
MB100G
EQ100A
EQ100G
FB100A
FB100G
MB20A
MB20G
EQ20A
EQ20G
FB20A
FB20G
Treatment

**Figure S3, A, B, C. Individual replicate population figures for post mating fecundity, of females from the sex ratio and diet regimes exposed for 3 days prior to mating to conspecifics or left alone.** Females from the Female-biased (FB), Equal sex (EQ) or Male –biased (MB) sex ratios and standard protein (High) or 20% protein (Low) diet regimes were tested. Females were either housed alone (A) or grouped with three rivals (G) for 3 days prior to mating. All conspecific non focal females and males were standard wild type.  Boxplots as per Fig.S1.

A

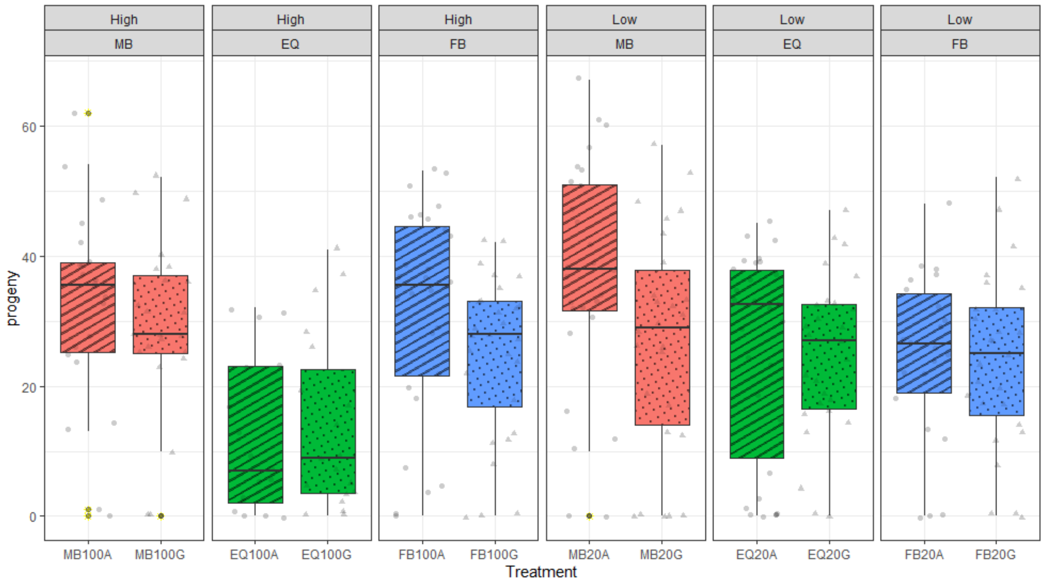
High
MB
High
EQ
High
FB
Low
MB
Low
EQ
Low
FB
progeny
0
20
40
60
MB100A
MB100G
EQ100A
EQ100G
FB100A
FB100G
MB20A
MB20G
EQ20A
EQ20G
FB20A
FB20G
Treatment

B

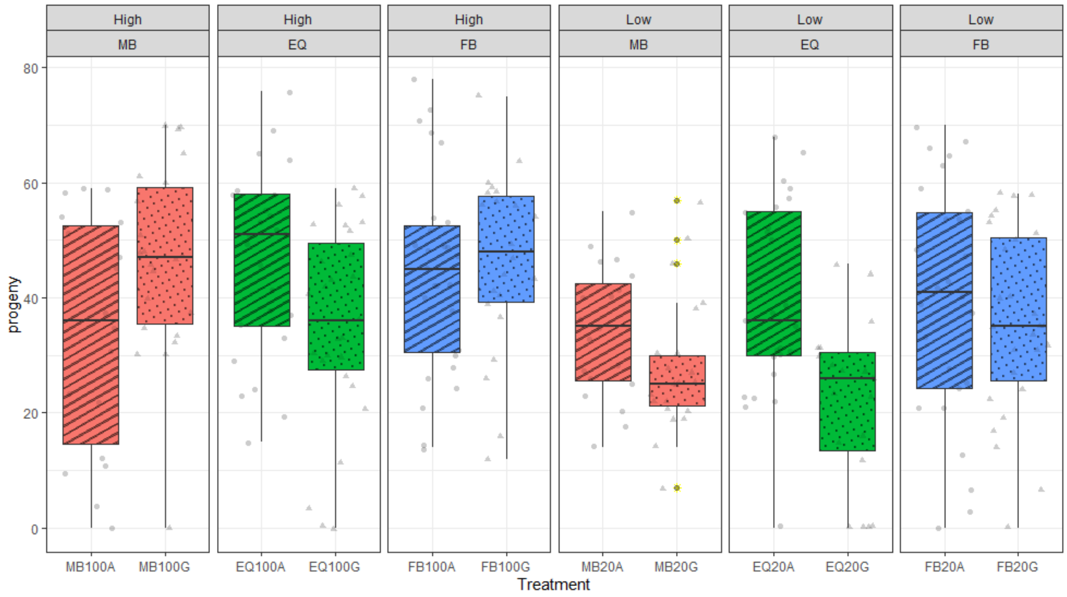
High
MB
High
EQ
High
FB
Low
MB
Low
EQ
Low
FB
progeny
0
20
40
60
80
MB100A
MB100G
EQ100A
EQ100G
FB100A
FB100G
MB20A
MB20G
EQ20A
EQ20G
FB20A
FB20G
Treatment

C

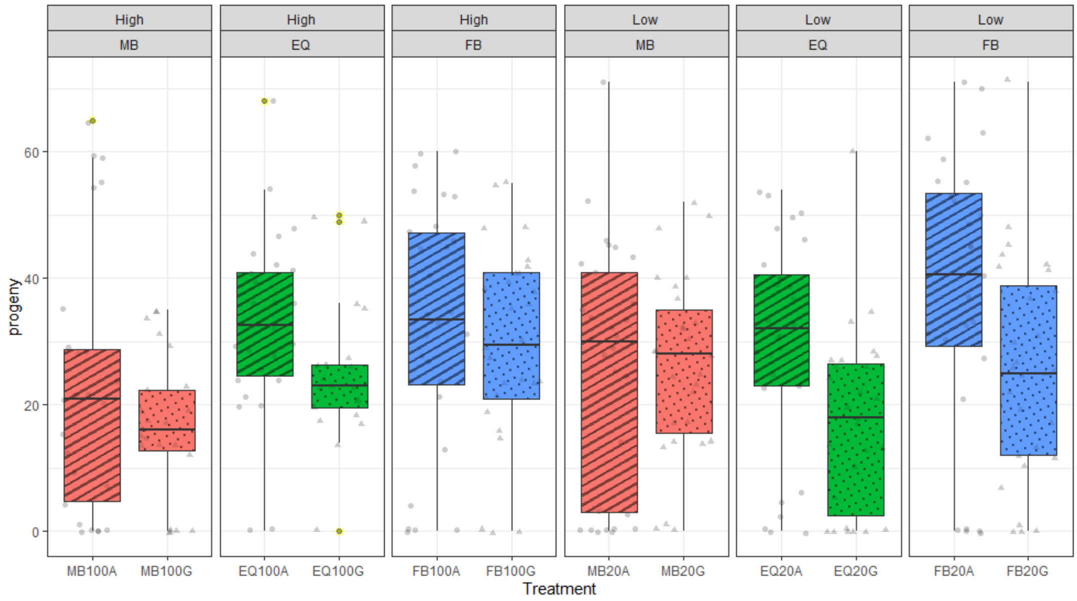
High
MB
High
EQ
High
FB
Low
MB
Low
EQ
Low
FB
progeny
0
20
40
60
MB100A
MB100G
EQ100A
EQ100G
FB100A
FB100G
MB20A
MB20G
EQ20A
EQ20G
FB20A
FB20G
Treatment

**Figure S4, A, B, C Individual replicate population progeny production (number of progeny emerging from the eggs laid in per female per 24h following mating), of females from the sex ratio and diet regimes exposed for 3 days prior to mating to conspecifics or left alone.** Females from the Female-biased (FB), Equal sex (EQ) or Male –biased (MB) sex ratios and standard protein (High) or 20% protein (Low) diet regimes were tested. Experimental females were either housed alone (A) or grouped with three rivals (G) prior to mating assay. All conspecific non focal females and males were standard wild type. Boxplots as per Fig.S1.

A

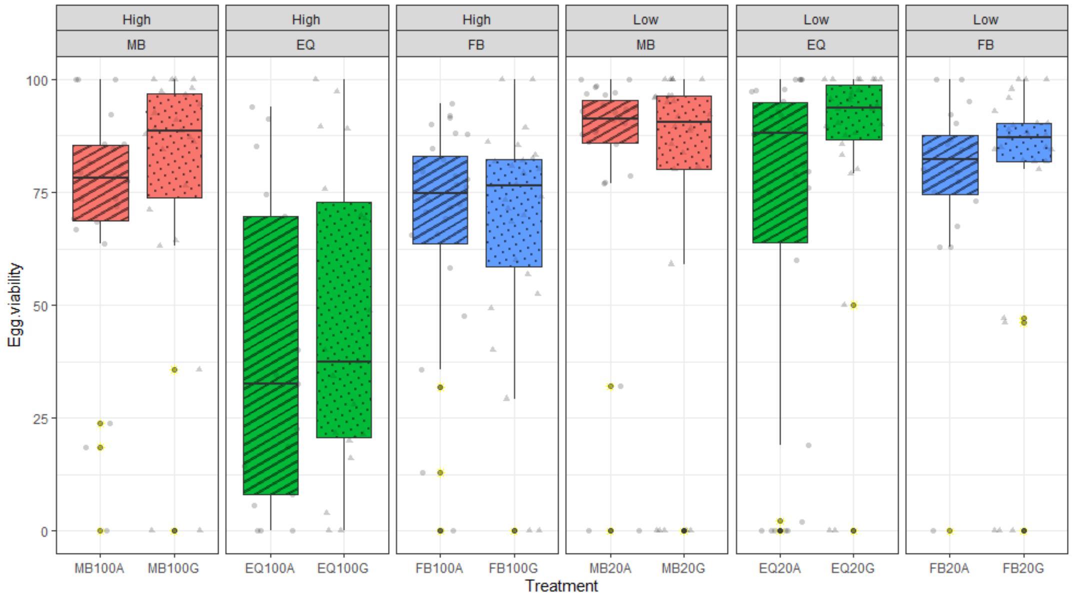
High
MB
High
EQ
High
FB
Low
MB
Low
EQ
Low
FB
Egg.viability
100
75
50
25
0
MB100A
MB100G
EQ100A
EQ100G
FB100A
FB100G
MB20A
MB20G
EQ20A
EQ20G
FB20A
FB20G
Treatment

B

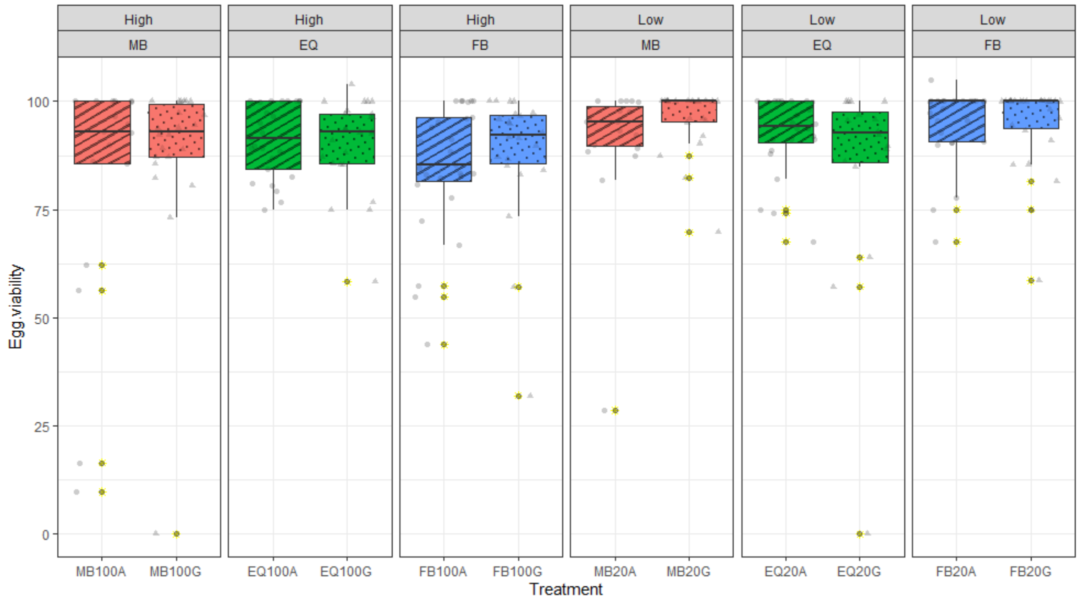
High
MB
High
EQ
High
FB
Low
MB
Low
EQ
Low
FB
Egg.viability
100
75
50
25
0
MB100A
MB100G
EQ100A
EQ100G
FB100A
FB100G
MB20A
MB20G
EQ20A
EQ20G
FB20A
FB20G
Treatment

C

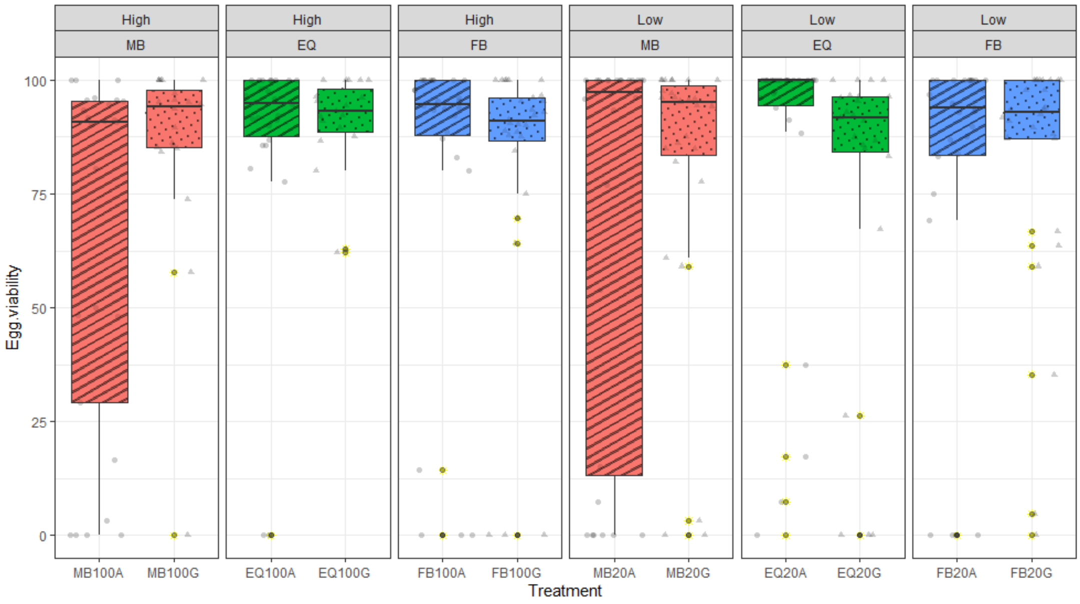
High
MB
High
EQ
High
FB
Low
MB
Low
EQ
Low
FB
Egg.viability
100
75
50
25
0
MB100A
MB100G
EQ100A
EQ100G
FB100A
FB100G
MB20A
MB20G
EQ20A
EQ20G
FB20A
FB20G
Treatment

**Figure S5, A, B, C. Individual replicate population Egg to adult viability (number of progeny / eggs per female per 24h following mating) of females from the sex ratio and diet regimes exposed for 3 days prior to mating to conspecifics or left alone.** Females from the Female-biased (FB), Equal sex (EQ) or Male –biased (MB) sex ratios and standard protein (High) or 20% protein (Low) diet regimes were tested. Experimental females were either housed alone (A) or grouped with three rivals (G) prior to mating assay. All conspecific non focal females and males were standard wild type. Boxplots as per Fig.S1.

**Table S1. Model results for analysis of number of eggs laid per 24 h after mating according to sex ratio (A) and diet (B) treatments.** Females from the Female-biased (FB), Equal sex (EQ) or Male –biased (MB) sex ratios and standard protein (High) or 20% protein (Low) diet regimes were tested. Experimental females were either housed alone or grouped with three rivals prior to mating assay. All conspecific non focal females and males were standard wild type.

**A – Effect of evolutionary sex ratio regime**

| Contrast | Estimate | SE | DF | t.ratio | p.value |
|---|---|---|---|---|---|
| EQ-FB | -6.52 | 4.22 | 11.9 | -1.544 | 0.3064 |
| EQ- MB | -3.68 | 4.24 | 12.1 | -0.868 | 0.6695 |
| FB-MB | 2.84 | 4.22 | 12.0 | 0.673 | 0.7813 |

**B – Effect of evolutionary diet regime**

| Contrast | Estimate | SE | DF | t.ratio | p.value |
|---|---|---|---|---|---|
| High – Low | 5.23 | 3.45 | 12 | 1.515 | 0.1558 |

**Table S2: Model results for analysis of egg to adult viability differences according to sex ratio (A), diet (B) or social (C) treatment effects.** Females from the Female-biased (FB), Equal sex (EQ) or Male –biased (MB) sex ratios and standard protein (High) or 20% protein (Low) diet regimes were tested. Experimental females were either housed alone or grouped with three rivals prior to mating assay. All conspecific non focal females and males were standard wild type.

**A – Effect of evolutionary sex ratio regime**

| Contrast | Estimate | SE | DF | t.ratio | p.value |
|---|---|---|---|---|---|
| EQ-FB | -3.02 | 7.81 | 12 | -0.386 | 0.9216 |
| EQ-MB | -1.89 | 7.83 | 12.1 | -0.241 | 0.9686 |
| FB-MB | 1.13 | 7.81 | 11.9 | 0.145 | 0.9885 |

**B – Effect of evolutionary diet regime**

| Contrast | Estimate | SE | DF | t.ratio | p.value |
|---|---|---|---|---|---|
| High – Low | -6.2 | 6.38 | 12 | -0.971 | 0.3506 |

**C – Effect of social treatment (Alone versus Grouped)**

| Contrast | Estimate | SE | DF | t.ratio | p.value |
|---|---|---|---|---|---|
| Alone – Grouped | -3.39 | 1.86 | 842 | -1.818 | 0.0694 |

**Table S3: Scripts of final LMER models (after model simplification) that were run using the lme4 package, on the response variables of eggs, progeny and mating latency.** Females from the Female-biased (FB), Equal sex (EQ) or Male –biased (MB) (**Sex.Ratio**) and standard protein (High) or 20% protein (Low) diet regimes were tested (**Food**). Experimental females were either housed Alone (A) or Grouped (G) with three rivals prior to mating assay (**Social**). All conspecific non focal females and males were standard wild type. Replicates identified as **Population.**

| Final Simplified Model | DF | Treatment | P value |
|---|---|---|---|
| Eggs ~ Sex.Ratio*Social*Food + (1\|Population) | 14 | Social (A vs G) | $6.49 \times 10^{-3}$ |
| Progeny ~ Sex.Ratio*Social*Food + (1\|Population) | 14 | Social (A vs G) | 0.0232 |
| LogLatency ~ Social + (1\|Population) | 1 | Social (A vs G) | 0.153 |

**Table S4: Script of final GLM model that was run using the lme4 package, on the response variable of non egg latying virguin females.** Females from the Female-biased (FB), Equal sex (EQ) or Male –biased (MB) (**Sex.Ratio**) and standard protein (High) or 20% protein (Low) diet regimes were tested (**Food)** for number of non egg laying females (**Zero Counts**). Experimental females were either housed alone (A) or grouped (G) with three rivals prior to mating assay (**Social**). All conspecific non focal females and males were standard wild type.

| **Model** | **DF** | **Factor** | **P** |
|---|---|---|---|
| Zero Counts ~ Sex.Ratio+Social+Food +Sex.Ratio:Food | 7 | Social | $6.44 \times 10^{-16}$ |
| | | Sex Ratio.MB.FoodLow | 0.22143 |
